# Emotional Modulation of Face Working Memory Precision in Specific Learning Disorders: Valence Flattening and Mnemonic Distortions

**DOI:** 10.64898/2026.09.13.751264

**Authors:** Marjan Makhsous, Mohsen Honar, Shaid Akbari, Ehsan Rezayat

**Author notes:** Corresponding Author: Ehsan Rezayat Department of Cognitive Sciences, Faculty of Psychology and Education, University of Tehran, Tehran, Iran.

## Abstract

This study examined whether emotional valence modulates face working memory precision in adolescents with specific learning disorders (SLD) and whether mnemonic distortions differ across subtypes. Fifty-four adolescents with SLD (dyslexia: n = 19; dysgraphia: n = 18; dyscalculia: n = 17) and 42 typically developing peers completed a morphed delayed reproduction task using sad, neutral, and happy faces. Using a von Mises + uniform mixture model, we decomposed recall errors into precision, systematic bias, and guessing. The SLD group made significantly larger errors (r = −0.61) and responded more slowly (r = −0.33). Mixture modeling showed that deficits arose from reduced precision (κ: TD M = 8.72 vs. SLD M = 4.15; r = 0.54) and stronger systematic bias toward neutrality (μ: TD M = −0.66° vs. SLD M = −4.15°; r = 0.34), with no differences in random guessing. Valence flattening was consistently larger in SLD. Among subtypes, dyscalculia showed the greatest working memory impairment, yet all three exhibited the same valence flattening pattern. Thus, emotional faces disrupt face working memory in SLD through reduced precision and systematic affective distortion. The bias parameter from mixture modeling may offer a clinically useful metric for subtype-sensitive assessment.

## INTRODUCTION

Specific learning disorders (SLDs)—including dyslexia, dyscalculia, and dysgraphia—are neurodevelopmental conditions that persist despite adequate intelligence and educational opportunity (American Psychiatric Association, 2013). While their primary manifestations are academic, accumulating evidence indicates that SLDs involve broader cognitive and affective disruptions extending beyond the classroom.

A lot of students with SLD don’t just struggle in academic settings; many of the same difficulties show up in everyday thinking as well. Recent research has made it clearer that these challenges are broader than people often assume. They can involve slower processing, weaker attentional control, and—maybe the most central part—problems with working memory (Daniel et al., 2022). When these systems are under pressure, it changes how a person handles information in the moment. Sometimes it’s something simple, like trying to follow a short instruction. Other times it’s more social—missing small cues in a conversation or losing track of what someone just said. So the impact isn’t limited to the classroom at all; it spills over into daily life.

Working memory — the ability to hold and manipulate information over brief periods — is at the core of these limitations (Peng & Fuchs, 2016). Verbal working memory issues, especially phonological loop problems in dyslexia, are well known. But newer research suggests that the nonverbal and perceptual sides of working memory are involved as well (Bozkurt et al., 2024). In everyday tasks, students with SLD often find it hard to pick out small visual details, and their eye-movement patterns can look a bit different when they’re trying to understand a crowded or complex scene. They also tend to take longer to encode things that change or move, compared with other students (Operto et al., 2020; Wang et al., 2025). Faces make things even more complicated. They carry emotional signals, but understanding those signals also requires attention and executive control, and that combination can be genuinely difficult for students with SLD. Many of them show weaknesses in both emotion recognition and working memory (Cromheeke & Mueller, 2016; Operto et al., 2020). Emotional expressions don’t always land the way they should—sometimes a sad face is interpreted as neutral (Bozkurt et al., 2024). Anxiety, which is common in SLD because of school pressure, drains the cognitive resources that should be helping with the task (Eysenck & Calvo, 1992). And when arousal rises too much, it throws off the balance between emotion and executive control, pulling working memory away from the information that actually matters (Teoh et al., 2024).

This is where the choice of task becomes important. Traditional match-to-sample tasks give you a simple right or wrong answer. They’re fine, but they miss the subtler distortions. Delayed reproduction tasks with morphed faces give you a continuous measure — not just whether someone remembered the face, but how accurately and in which direction they misremembered it (Bays et al., 2009). That directionality — a systematic drift toward neutrality, what we call valence flattening — opens a window into executive affective filtering failures (Authors et al., 2025). And with mixture models, you can break down errors into separate components: precision, bias, and random guessing. That’s the kind of detail you just don’t get with span-based assessments.

Still, one question has remained unanswered: how do these emotion-driven distortions play out across different SLD subtypes? We know that subtypes have distinct cognitive profiles. For example, dyscalculia is often linked to weaknesses in visuospatial working memory (David, 2012), whereas dyslexia tends to involve phonological and perceptual encoding difficulties (Daniel et al., 2022). Even though these differences are well established, no study has directly compared how emotional valence shapes face-based working memory errors — using both precision and bias — across dyslexia, dyscalculia, and dysgraphia, alongside typically developing peers.

So, we designed the present study to fill that gap. We used a morph-based delayed reproduction task and decomposed recall errors — via mixture modeling — into precision (κ), bias (μ), and guessing (γ). Our main aim was to see whether emotional valence affects mnemonic precision differently across SLD subgroups, and whether valence flattening turns out to be a general feature of SLD or something subtype-specific. In doing so, we hope to clarify how affective load contributes to SLD and to move toward more sensitive, subtype-aware diagnostic and intervention strategies.

## METHODS

### Participants

We started with 110 adolescents. After excluding 14 who couldn’t complete both tasks or had missing data across conditions, we ended up with a final sample of 96 — 42 typically developing (TD) adolescents and 54 with a confirmed diagnosis of specific learning disorder (SLD). Diagnoses were made by a licensed physician or clinical psychologist, and we double-checked using structured parent interviews and the Colorado Learning Disabilities Questionnaire (CLDQ). Within the SLD group, 19 had dyslexia, 18 had dysgraphia, and 17 had dyscalculia. A few participants with mixed or unspecified profiles stayed in the overall SLD group but weren’t included in the subtype-specific analyses.

All parents or legal guardians gave written informed consent before the study began. The Institutional Ethics Committee of the University of Tehran approved the procedures (Protocol ID: IR.UT.PSYEDU.REC.1404.102), and we followed the Declaration of Helsinki and its later updates. Since we were working with adolescents, we made sure every step was carried out with care and sensitivity.

### Stimuli and Apparatus

To test emotional perception and face working memory, we created 19-step morph continua starting from a gender-neutral, emotion-neutral base face, using FaceGen Modeller 3.5 and InterFace software (Kramer et al., 2017). Each step changed the expression by 10%, giving us images ranging from 90% sad to 90% happy (see Figure 1). For the actual analyses, we picked three images per emotion: indices 2, 4, and 6 for sad; 8, 10, and 12 for neutral; and 14, 16, and 18 for happy. All faces were cropped into ovals in Photoshop and matched for luminance using the SHINE toolbox in MATLAB (Willenbockel et al., 2010). They subtended about 3.33° × 4.76° of visual angle and were shown in the centre of a 23-inch TFT monitor (1920 × 1080 px, 60 Hz). The whole experiment ran in MATLAB R2019b with PsychToolbox-3.

**Figure 1.**
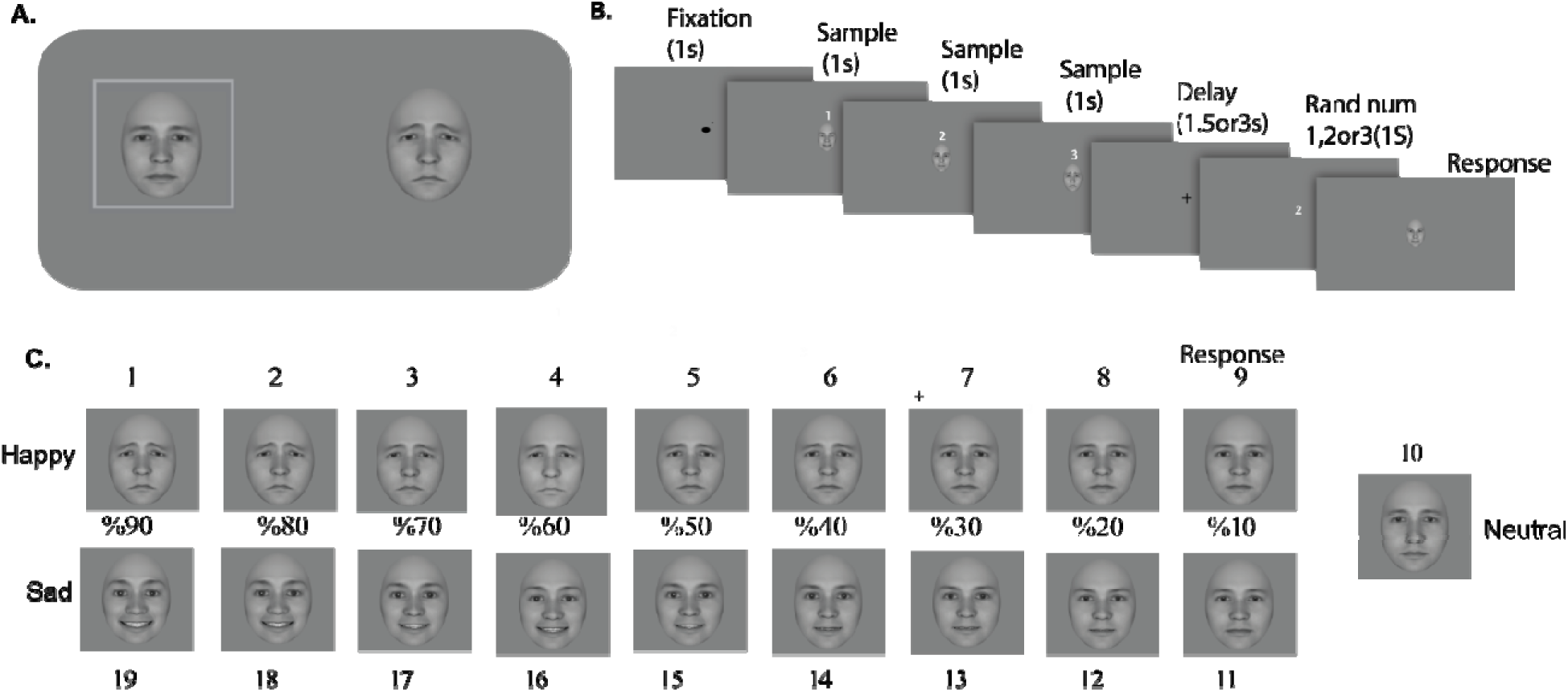
Experimental procedure and stimuli. (A) Perceptual matching task: participants adjusted a probe face (right) to match a reference face (left) using arrow keys. This task measured perceptual ability without memory load. (B) Delayed-reproduction trial sequence: after a 1-s fixation, three sample faces appeared sequentially (1 s each), followed by a 1.5- or 3-s delay. A numeric cue then prompted recall of one of the three faces, and participants adjusted a test face to match the target. (C) Morph continuum: 19 graded images ranging from 90% sad (left) to 90% happy (right), generated in 10% steps from a neutral base face.

### Procedure

Participants sat in a dim, quiet room about 60 cm from the screen. After five practice runs, they completed two tasks, always in the same order.

#### Perceptual Matching Task

Two faces appeared side by side — a fixed reference on the left and an adjustable probe on the right. Using arrow keys, participants morphed the probe until it matched the reference’s emotional intensity along the 19-step scale, then hit SPACE to confirm. This gave us a baseline measure of perceptual ability, without any memory load.

#### Face Working Memory Task (Delayed Reproduction)

Each trial started with a 1-second fixation point. Then three sample faces appeared one after the other (1 second each), labelled 1 to 3, randomly picked from a set of morph levels {0, 20, 40, 60, 80%}. After a delay of either 1.5 or 3 seconds, a number appeared, telling participants which face to recall. An adjustable test face then came up, and they used arrow keys to match it to the remembered target, pressing SPACE to submit their answer. Everyone completed three blocks of 54 trials each (2 delays × 9 stimuli × 3 repetitions), with 3-to 5-minute breaks in between. To prevent lucky guesses, we made sure the distractor faces were always at least six morph steps away from the correct target.

### Data Analysis

#### Preprocessing

For both tasks, we removed trials where responses were missing (reaction time = 0 or response value = 0). Any response above the valid morph range (> 19) was corrected using circular fold-back (abs(response − 38)), because the morph continuum is circular. This affected fewer than 2% of all trials.

#### Error Metrics

Absolute error was simply the difference between the response and the target, ignoring direction: Error = |Response − Target|. These values ranged from 0 to 18 and gave us a clean measure of recall precision. Signed error was Response − Target, with positive values meaning a drift toward happier expressions and negative values meaning a drift toward sadder ones. This became our primary index of directional bias — our marker for valence flattening.

#### Subject-Level Aggregation

All our statistical tests were done at the subject level. For each participant, we averaged absolute error, signed error, and reaction time separately for sad, neutral, and happy conditions, and for the two delay lengths (1.5 s and 3 s) in the FWM task. This treats each person as a single independent observation and avoids the inflated sample sizes that come from pooling trial-level data.

Delay conditions (1.5 s and 3 s) were collapsed for primary analyses, as preliminary comparisons revealed no significant interaction between delay length and group across either task (all p > .12).

#### Nonparametric Statistical Tests

Histograms and Q-Q plots told us the error distributions weren’t normal, and Shapiro-Wilk tests confirmed it. So we stuck with nonparametric methods throughout. Between-group comparisons (TD vs. SLD) used two-tailed Mann-Whitney U tests. Effect sizes were calculated as rank-biserial correlations (r_rb = (U□− U□) / (n□× n□)), with .10 = small, .30 = medium, and .50 = large (Kerby, 2014). For subgroup comparisons across four groups (TD, dyslexia, dysgraphia, dyscalculia), we ran Kruskal-Wallis tests first, and when the omnibus effect was significant, we followed up with Mann-Whitney U tests and Bonferroni corrections (alpha = .05/6 = .0083). Within-group signed error analyses used one-sample Wilcoxon tests against zero, with Bonferroni adjustments across the three emotion conditions (alpha = .05/3 = .017).

#### Mixture Modeling of Memory Errors

To break down recall errors into meaningful cognitive components, we fitted a von Mises + uniform mixture model to each participant’s signed error distribution, separately for each emotion and overall (Bays et al., 2009). The model assumes that responses come from one of two processes: memory-based responses (a von Mises distribution with mean μ and concentration κ) or random guesses (a uniform distribution with weight γ). Higher κ means sharper, more precise memory (Authors et al., 2024); μ tells us about systematic drift; and γ tells us how often the participant is just guessing.

We estimated parameters using expectation-maximization (EM) with a strict convergence criterion (||Δparams|| < 10□□) and up to 2,000 iterations. Fits that didn’t converge (less than 1% of cases) were excluded. Group comparisons on μ, κ, and γ used Mann-Whitney U tests; subgroup comparisons used Kruskal-Wallis followed by Bonferroni-corrected post-hoc tests, just like before. The bias parameter μ (in degrees, where 10° is roughly one morph step) directly measures valence flattening: positive μ for sad faces means drift toward neutrality; negative μ for happy faces also means drift toward neutrality. Precision (κ) captures mnemonic fidelity, and guess rate (γ) indexes how often participants disengage from the task and respond randomly.

## RESULTS

Our final sample included 42 typically developing adolescents (18 boys, 24 girls; age range 10– 16, mean = 12.7, SD = 1.72) and 54 adolescents with SLD (27 boys, 27 girls; age range 10–15, mean = 11.8, SD = 1.50). Table 1 gives the full demographic breakdown.

**Table 1:** Demographic Characteristics of Participants.

| Variable | TD (n = 42) | SLD (n = 54) | Total (N = 96) |
| --- | --- | --- | --- |
| Age (years), M (SD) | 12.7 (1.72) | 11.8 (1.50) | — |
| Age Range (years) | 10–16 | 10–15 | 10–16 |
| **Sex, n (%)** |  |  |  |
| Boys | 18 (42.9%) | 27 (50.0%) | 45 (46.9%) |
| Girls | 24 (57.1%) | 27 (50.0%) | 51 (53.1%) |
| **SLD Subtype, n (%)** |  |  |  |
| Dyslexia | — | 19 (35.2%) | 19 (19.8%) |
| Dysgraphia | — | 18 (33.3%) | 18 (18.8%) |
| Dyscalculia | — | 17 (31.5%) | 17 (17.7%) |
| Unspecified/Comorbid | — | Excluded from subgroup analyses |  |

*Note.* TD = Typically Developing; SLD = Specific Learning Disorder. Age reported as mean (SD). Participants with comorbid or unspecified SLD profiles were included in overall SLD analyses but excluded from subgroup comparisons.

Let’s start with the big picture. The SLD group made significantly more errors than the TD group in both tasks. In the Face Working Memory task, the difference was substantial (z = −5.07, p < .001, rank-biserial r = −0.61). In the Perception task, the effect was smaller and just shy of significance (z = −1.92, p = .055, r = −0.23). Still, the overall direction was consistent: SLD adolescents struggled more with faces, whether they had to remember them or just match them (Figure 2). Neither group showed a significant difference between the two tasks, so we’re not looking at a task-specific effect — it’s more general.

**Figure 2.**
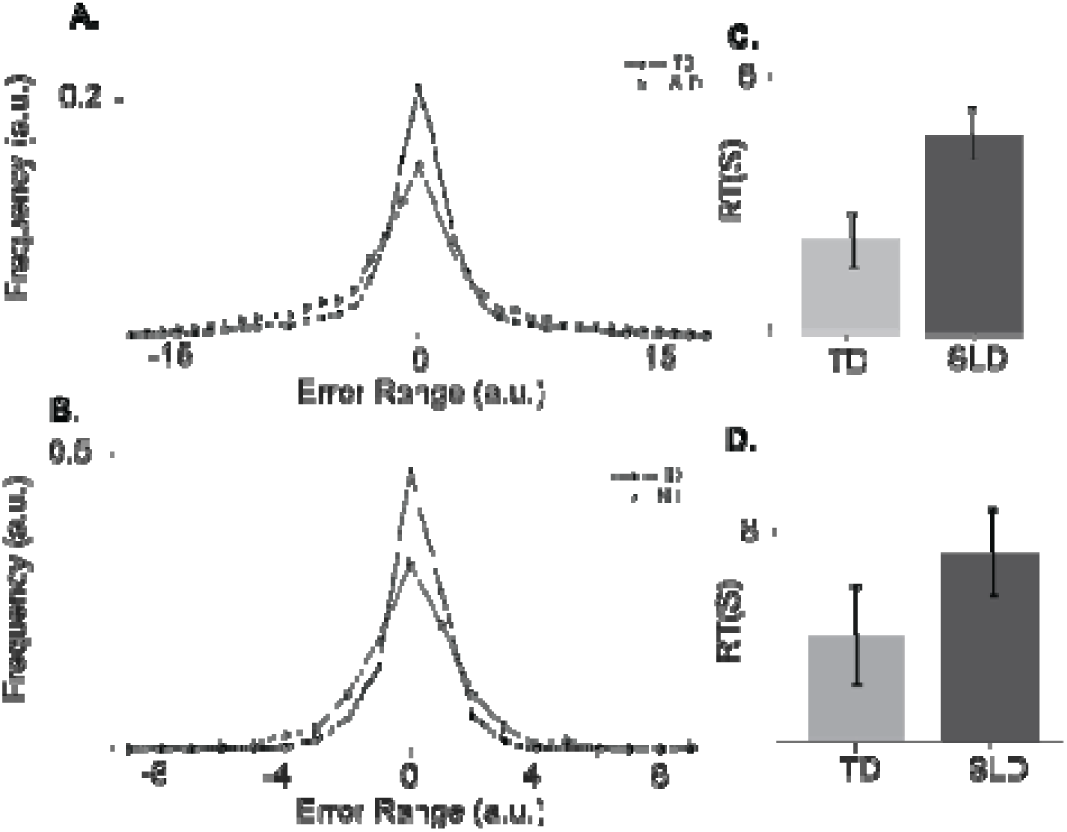
Group differences in accuracy and reaction time. (A) Face Working Memory accuracy (absolute error): SLD participants made significantly larger errors than TD controls (*p* < .001, *r* = −0.61). (B) Perception task accuracy: SLD participants showed a trend toward larger errors (*p* = .055, *r* = −0.23). (C) Reaction times in the Face WM task: SLD participants were slower (*p* = .006, *r* = −0.33). (D) Reaction times in the Perception task: SLD participants were also slower, though the difference did not reach conventional significance (*p* = .076, *r* = −0.22). Overall, the pattern suggests slower and less accurate performance in SLD across both tasks.

Reaction times told a clearer story. In the Face WM task, SLD participants were noticeably slower (z = −2.74, p = .006, r = −0.33). In the Perception task, they were also slower, though the difference didn’t quite reach conventional significance (z = −1.78, p = .076, r = −0.22). But the pattern is consistent: SLD adolescents aren’t rushing — they’re taking longer, which fits with the idea of processing efficiency limitations rather than impulsivity (Figures 2C-D).

Now, what about specific emotions? Across the board, TD participants outperformed SLD participants, but the gap wasn’t equal for all emotions. The largest difference showed up for happy faces, where the SLD group really struggled. The numbers: for sad faces, z = −3.70, p < .001, r = −0.44; for neutral, z = −4.22, p < .001, r = −0.50; and for happy, z = −5.33, p < .001, r = −0.64 (Figure 3). Within the SLD group, sad faces actually yielded the best accuracy, but the differences across emotions weren’t significant — so the pattern isn’t that one emotion is “easier” for them; it’s more than happy faces are particularly hard.

**Figure 3.**
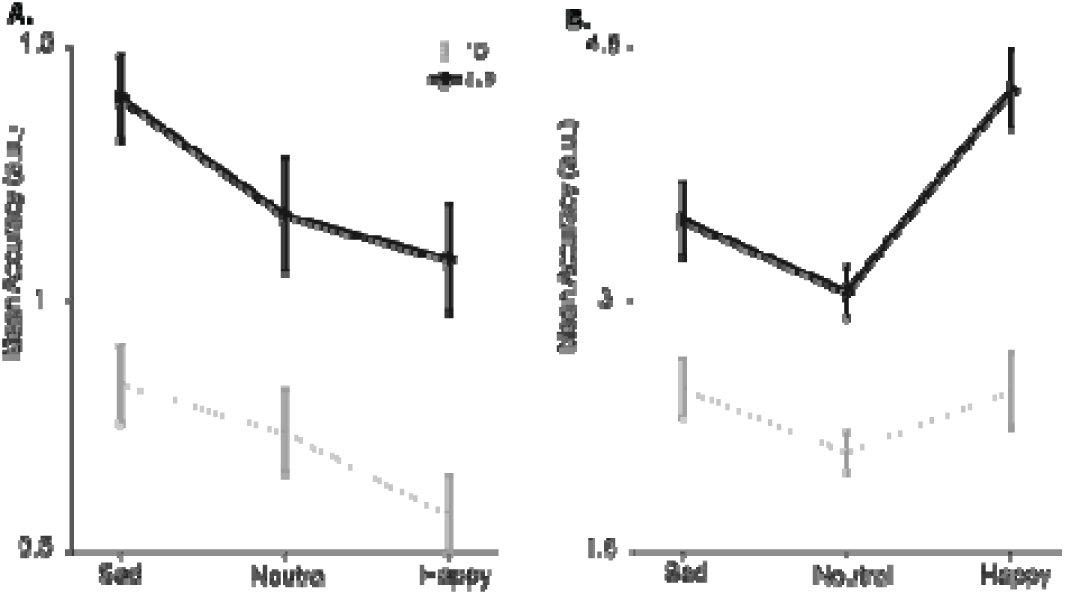
Emotion-specific accuracy across groups. (A) Perception task: TD participants outperformed SLD participants across all emotions, with the largest gap for happy faces. (B) Face Working Memory task: TD participants were more accurate across all emotions, with significant group differences for all three valence conditions (Sad: *z* = −3.70, *p* < .001, *r* = −0.44; Neutral: *z* = −4.22, *p* < .001, *r* = −0.50; Happy: *z* = −5.33, *p* < .001, *r* = −0.64), and the largest disparity emerging for happy faces. Within the SLD group, sad faces yielded the best accuracy, though differences across emotions were not significant. These results indicate that happy faces are particularly challenging for SLD adolescents, both in perception and in memory.

When we broke things down by subtype, the picture got more interesting. A Kruskal-Wallis test showed significant differences across the four groups (H = 29.12, p < .001). All three SLD subtypes performed worse than TD controls, but the order mattered. Dyscalculia showed the largest working memory errors (mean = 4.06 morph steps), followed by dyslexia (3.66) and dysgraphia (3.17). All three were significantly behind TD, with effect sizes ranging from r = −0.46 to −0.76 (Figure 4). That’s a wide spread, and it tells us that not all SLD subtypes are equally affected when it comes to face working memory.

**Figure 4.**
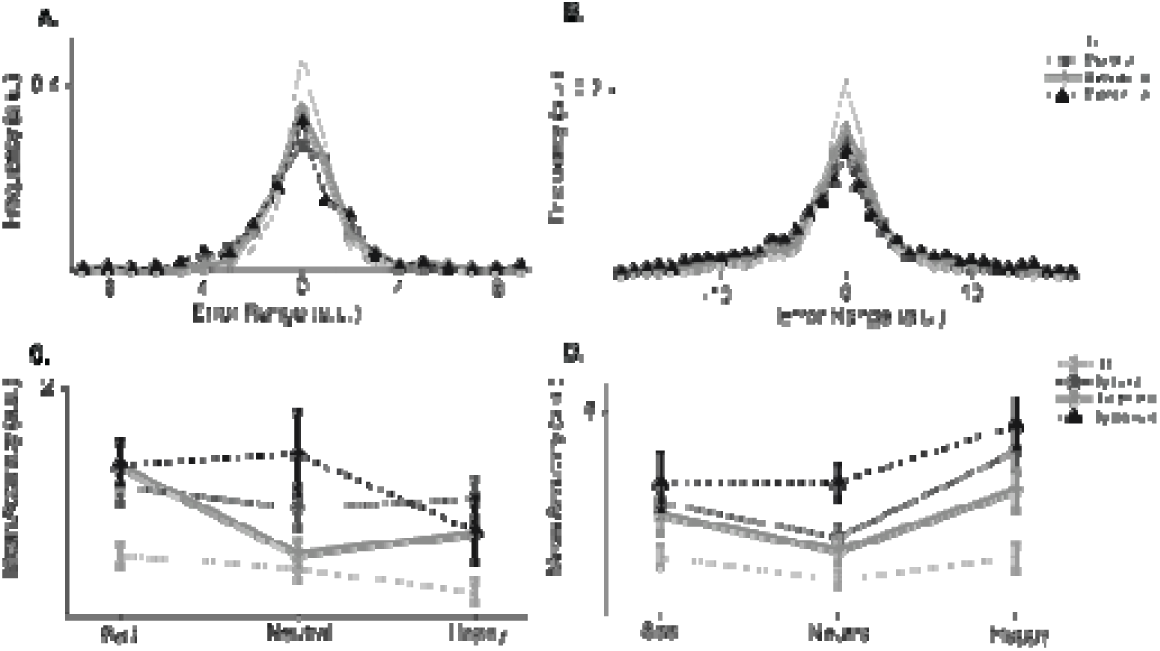
Subgroup differences in task performance. (A) Perception task baseline: all three SLD subtypes showed numerically larger errors than TD, with dyslexia showing the highest values, though no subgroup differences reached significance overall (Kruskal-Wallis *p* = .235). (B) Face Working Memory baseline: significant group differences emerged (Kruskal-Wallis *p* < .001). Dyscalculia showed the largest errors (mean = 4.06 morph steps), followed by dyslexia (3.66) and dysgraphia (3.17). All three were significantly behind TD. (C) Perception task with emotional modulation: TD and dysgraphia showed relatively stable performance across emotions, while dyslexia showed more variability. (D) Face Working Memory with emotional modulation: emotional valence affected all groups, but the impact was largest in dyslexia and dyscalculia. These findings point to distinct cognitive profiles across SLD subtypes — perceptual encoding difficulties in dyslexia, and visuospatial maintenance difficulties in dyscalculia.

The mixture model provided a clearer picture of what was driving these group differences (Figure 5). Memory precision (κ) was significantly lower in the SLD group than in TD controls across all three emotional conditions. For sad faces, TD participants averaged 9.45 compared to 5.85 for SLD (*z* = 3.65, *p* < .001, *r* = 0.44); for neutral faces, the gap was similar (TD M = 10.75 vs. SLD M = 4.80; *z* = 3.74, *p* < .001, *r* = 0.45); and for happy faces — where the difference was largest — TD averaged 15.05 while SLD dropped to 6.54 (*z* = 4.60, *p* < .001, *r* = 0.55). Crucially, guess rates (γ) did not differ significantly between groups under any emotion condition (all *p* > .35), indicating that SLD adolescents were actively attempting retrieval rather than responding randomly — their memory representations were simply less precise.

**Figure 5.**
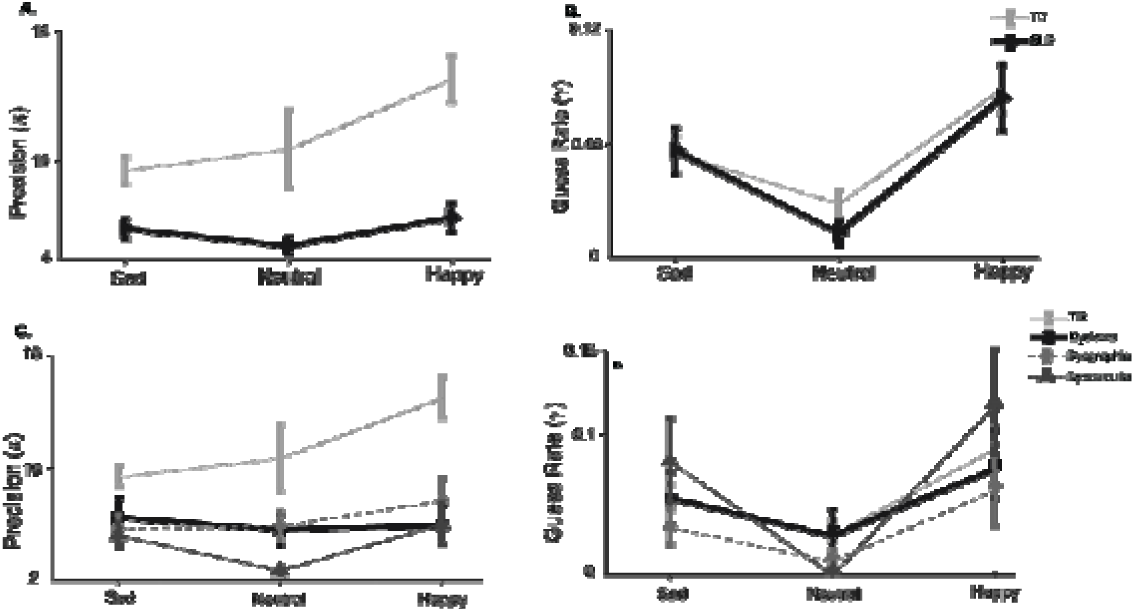
Mixture model parameters across groups and emotions. (A) Memory precision (κ) for TD and SLD groups: TD participants showed substantially higher precision across all emotional conditions, with both groups peaking for happy faces. (B) Guess rate (γ) for TD and SLD groups: guess rates were low and comparable between groups across all conditions, confirming that SLD deficits reflect distorted rather than absent memory representations. (C) Precision (κ) across SLD subtypes: all three subtypes showed markedly lower precision than TD controls, with Dyscalculia showing the lowest values particularly for Neutral faces. (D) Guess rate (γ) across SLD subtypes: guess rates were generally low across all subtypes, though Dyscalculia showed elevated guessing for Sad faces. Error bars represent ±1 SEM.

Subgroup analyses of precision revealed significant Kruskal-Wallis effects for all emotion conditions (all *p* < .004), with each of the three SLD subtypes showing lower κ than TD controls. Dyscalculia showed the greatest deficit overall, particularly for neutral faces (*z* = 4.43, *p* < .001, *r* = 0.74). Guess rates remained comparably low across subtypes, with Dyscalculia showing a slight elevation in random responding for sad faces relative to the other groups.

Now, the valence flattening effect. Both groups showed a systematic drift toward neutrality — that is, when remembering sad faces, they tended to report them as less sad; when remembering happy faces, they reported them as less happy. But the magnitude was much larger in the SLD group. For sad faces, TD drifted by +1.67 morph steps on average, while SLD drifted by +2.70. For neutral faces, TD drifted by −0.34, SLD by −0.73. And for happy faces — the most dramatic difference — TD drifted by −1.75, while SLD drifted by −3.68. All of these differences were significant (all p < .05; Figure 6). The SLD group also showed wider error distributions overall, especially for happy faces, where the gap between groups was largest.

**Figure 6.**
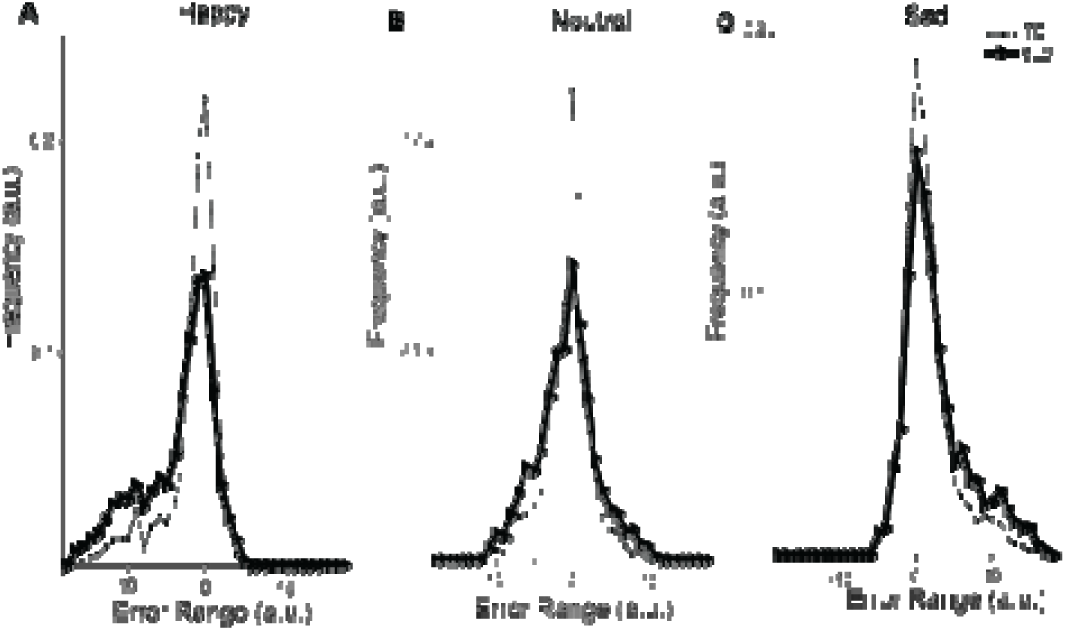
Valence flattening: systematic drift toward neutrality. Distributions of signed errors for (A) Happy, (B) Neutral, and (C) Sad faces, shown separately for TD (dotted line) and SLD (solid line) groups. Negative values indicate drift toward sadder expressions; positive values indicate drift toward happier expressions. Both groups showed a systematic drift toward neutrality for happy and sad faces — but the effect was consistently larger in the SLD group. For happy faces, SLD participants drifted by −3.68 morph steps on average, nearly twice the TD drift (−1.75). The SLD group also showed wider, more variable error distributions overall. This pattern confirms that valence flattening — a breakdown in executive-affective filtering — is present in both groups but significantly amplified in SLD.

So, putting it all together: adolescents with SLD showed clear impairments in both face memory and emotion perception. Their errors were larger, their responses were slower, and their memory traces were both less precise and more biased toward neutrality. Dyscalculia stood out as the most impaired subtype on working memory, but all three subtypes showed the same converging pattern of valence flattening. These results point to a fundamental disruption in how emotional information is maintained and retrieved — not a simple matter of effort or attention.

## DISCUSSION

So, what does all this actually tell us about how adolescents with SLD process and remember emotional faces? The broad strokes are clear enough — they’re slower and less accurate. But the real story is in the details.

Let’s start with what the mixture model showed. The SLD group didn’t just perform worse because they gave up or started guessing randomly. Their guess rates were statistically identical to the TD group. So, they were trying just as hard. The problem was that their memory representations were fuzzier (lower precision) and systematically pulled off-target (amplified bias). That’s a crucial distinction. It shifts the conversation from “SLD = poor memory” to a more precise question: how does memory fail? In this case, through reduced fidelity and affective distortion — not disengagement.

Now, here’s a finding that caught our attention. Happy faces weren’t just difficult for the SLD group — they were the most distorted. If you’d asked us before the study, we probably would have guessed that negative expressions like sadness would be more disruptive. But that’s not what we found. Why might that be? One possibility is that positive emotional content — a smiling face, for example — sets off a kind of cognitive friction. Maybe the brain has to adjust its expectations or shift internal states in a way that neutral or even sad faces just don’t trigger. For a group already dealing with executive control difficulties and heightened anxiety, this extra layer of demand might be enough to tip the system over.

This line of thinking fits with more recent work. It seems that positive valence isn’t always a good thing — especially when the executive system is already under pressure. For example, Provost et al. (2025) demonstrated that positive distractors impaired working memory accuracy under high cognitive load. Lautenbach (2024) came to a similar conclusion in an extensive meta-analysis, showing that positive emotions can have complex and even counterproductive effects on executive functions. But that’s not all. Neuroimaging studies tell the same story. Clarke and Johnstone (2013), for example, showed using fMRI that under high cognitive load, top-down control from the ventrolateral prefrontal cortex and dorsal anterior cingulate helps keep the amygdala in check and reduces emotional interference. Taken together, these findings suggest that when executive-affective filtering fails — as we observed in the SLD group — emotional distinctiveness is lost, and memory traces drift toward a blunted, neutral baseline.

The subtype differences are worth pausing over. Dyscalculia showed the most pronounced working memory impairment — the largest errors and the lowest precision across emotions. That makes sense if you think of dyscalculia as a disorder of visuospatial representation (David, 2012). Holding a complex face in mind, with all its subtle configural details, taps into the same visuospatial sketchpad that’s already compromised in this group. Dyslexia, on the other hand, showed a different profile. Its biggest deficits showed up in the perceptual task, especially for neutral faces. That fits with the idea that dyslexia involves a domain-general difficulty with categorical perception — a problem at the encoding stage, before information even reaches working memory (Daniel et al., 2022; Shaban et al., 2024). Dysgraphia, interestingly, consistently performed closest to the TD group. That’s consistent with its primary deficit being more motor-sequential than visuospatial or perceptual. So, while all three subgroups share a common thread — valence flattening — the route to that flattening appears to differ. Subgroup comparisons should be interpreted cautiously given the moderate subgroup sample sizes, and replicated in larger independent cohorts.

This is where the mixture model really earns its keep. By showing that guess rates are unchanged, we can rule out the idea that SLD adolescents are simply disengaging from the task. They’re not. They’re actively trying, but their memories are systematically distorted. That distinction has practical implications. In a classroom or clinic, a child who is working hard but producing biased responses needs a different kind of support than one who is randomly guessing. These findings suggest that interventions aimed at calibrating emotional representations — perhaps through feedback-based training or cognitive-reappraisal exercises — might be more effective than generic memory drills. On a more practical note, Authors et al. (2025) recently showed that even single-session interventions can modulate memory precision. That raises an intriguing possibility: brief, targeted training might help recalibrate this affective bias.

Of course, we have to be honest about where the picture is incomplete. Our design was cross-sectional, so we can’t talk about causality or development. We only used three emotions — sad, neutral, happy — leaving out fear, surprise, or disgust. And we only measured behaviour, so the neural mechanisms remain speculative. But these limitations also point the way forward. Incorporating EEG or fMRI could tell us whether the bias originates in early visual areas like the fusiform gyrus or later executive regions like the prefrontal cortex. Longitudinal studies could reveal whether valence flattening is stable or changes over time. And extending the paradigm to more naturalistic settings — using dynamic faces or real-time social interactions — would show whether these lab-based metrics actually translate to the playground or the classroom.

So, where does that leave us? The present work provides convergent evidence that emotional valence disrupts face working memory in SLD — not by making children forget, but by making their memories systematically less precise and more neutral. The dissociation across subtypes suggests that a one-size-fits-all approach won’t work. Instead, we need more tailored strategies: working on perceptual encoding in dyslexia, visuospatial maintenance in dyscalculia, and across the board, recalibrating how emotion and memory interact. Valence flattening — especially when measured through mixture-model bias parameters — seems to capture something fundamental about the intersection of emotion and cognition in SLD. Something that traditional measures simply miss. A formal a priori power analysis was not conducted; future studies should determine optimal sample sizes for subgroup comparisons.

